# Vector-mediated viral transmission favours less virulent viruses

**DOI:** 10.1101/323527

**Authors:** Emily J. Remnant, Niklas Mather, Thomas L. Gillard, Boris Yagound, Madeleine Beekman

**Affiliations:** Behaviour and Genetics of Social Insects Laboratory, School of Life and Environmental Sciences, The University of Sydney

**Keywords:** Honeybee viruses, virulence, evolution, vector transmission

## Abstract

While it is well-established that the ectoparasitic mite *Varroa destructor* is largely responsible for the widely-reported decline of populations of the Western honeybee *Apis mellifera*, the exact role the mite plays in honeybee health remains unclear. The last few years have seen a surge in studies associating RNA viruses vectored by the mite with the death of honeybee colonies. *Varroa* facilitates the spread of RNA viruses because it feeds on developing bee brood and transfers haemolymph from bee-to-bee. Such a change in transmission, from horizontal and vertical to vector-based, is predicted to lead to an increase in virulence of RNA viruses, thus potentially providing an explanation for the observed association between *Varroa* and certain viruses. Here we document the effect of changing the route of transmission of honeybee viruses contained in the haemolymph of honeybee pupae. We find that a change in mode of transmission rapidly increases viral titres of two honeybee viruses, Sacbrood virus (SBV) and Black queen cell virus (BQCV). This increase in viral titre is accompanied by an increase in virulence. In contrast, the virus most often associated with *Varroa*, Deformed wing virus (DWV), shows a reduction in viral titre in the presence of SBV and BQCV. In addition, DWV does not cause mortality to honeybee pupae in isolation. Most likely a change in mode of transmission due to the arrival of a vector quickly eliminates the most virulent honeybee viruses resulting in an association between *Varroa* and less virulent viruses such as DWV. Our work therefore provides empirical evidence for an alternative explanation for the widely-observed association between *Varroa* and DWV.

## Introduction

It is indisputable that the Western honeybee *Apis mellifera* suffers from the negative effects of inappropriate use of pesticides^1^ and a range of parasites and diseases^2^. The most important parasite today is the ectoparasitic mite *Varroa destructor*. The emergence of *V. destructor* is the result of a host shift that occurred when *A. mellifera* and the Asian Hive Bee, *A. cerana*, were brought into contact by beekeepers in the 1930s^3^.

*Varroa destructor* (hereafter simply referred to as *Varroa*) is aptly named. When left untreated, *Varroa* typically destroys the colonies of its host^4^. In Europe and the United States managed honeybee colonies suffer greatly from *Varroa* and require constant treatment with miticides to prevent colonies from dying. At the same time, wild or feral honeybee populations have been decimated or gone extinct^5^.

*Varroa* females feed on the haemolymph of the developing bees and in doing so are thought to vector viruses carried therein^6, 7^. Although a variety of viruses could potentially be transmitted by *Varroa^8^*, one in particular - Deformed wing virus (DWV)- is strongly associated with *Varroa*. For example, as *Varroa* sequentially invaded the islands of Hawaii, viral titres of DWV increased, while the diversity of DWV viral strains decreased, such that a single strain came to dominate after a few years^9^. A similar phenomenon was seen in New Zealand where titres of DWV dramatically increased with the length of exposure to Varroa^10^.

Vector-based transmission is predicted to lead to an increase in virulence because it changes the evolutionary trade-off between virulence and transmission^11^. While an obligate parasite is selected to replicate quickly, so that it can infect as many hosts as possible, a high rate of replication may kill the host before the parasite is transmitted to its next host. Selection will thus act against a pathogen that kills or immobilises its host if this reduces its long-term transmission success^12, 13^. The arrival of a vector changes the dynamics of the transmission-virulence trade-off. If a pathogen can harness a mobile vector to facilitate its spread to new hosts, then it no longer relies on its current host for transmission.

An increase in virulence after a change in route of transmission was recently documented in the obligate endosymbiont *Wolbachia* and one of its native hosts, the isopod *Armadillidium vulgare*. Because *Wolbachia* is normally transmitted vertically, via eggs, it requires its host to be alive and reach reproductive age. Hence, *Wolbachia* tends to form symbiotic relationships with its hosts. However, when the route of transmission was changed from vertical to horizontal, by injecting *Wolbachia* directly into the haemolymph of the host, *Wolbachia* titres quickly escalated and infections became highly virulent, resulting in the death of the hosts after only a few serial passages^14^.

At first glance the association between *Varroa* and DWV seems to fit the predicted change in virulence after the arrival of a vector, and thus a change in mode of transmission. However, honey bees host many viruses that are both common and widespread ^8,15^ including viruses that, like DWV, are present in *Varroa* and can also be vector-transmitted (eg. viruses of the Acute bee paralysis virus complex (ABPV and Kashmir bee virus, etc.)^16^). This raises the question: why has DWV become synonymous with *Varroa* infestation, but not other honey bee viruses? An alternative explanation for the observed association is that more virulent viruses are eliminated from the population due to excessive host mortality following vector-based transmission, thereby allowing less virulent DWV to take the upper hand^10, 17, 18^. Here we test this alternative explanation empirically using a population of honeybees naïve to both *Varroa* and DWV.

We experimentally changed the transmission of bee viruses from horizontal (via faeces and feeding) to vector-mediated transmission by performing a serial passage experiment. We injected extracts from bee pupa to bee pupa repeatedly for up to 30 transmission cycles and found that two viruses naturally present in our bee population, Sacbrood virus (SBV) and Black queen cell virus (BQCV), rapidly increased in titre. In contrast, DWV introduced via injection rapidly decreased in titre accompanied by a rapid increase in titres of SBV and BQCV. More importantly, DWV alone did not cause mortality in pupae, whereas injection with serially passaged bee extracts containing high titres of SBV and BQCV did. We conclude that the observed association between *Varroa* and DWV may not necessarily be due to *Varroa* increasing the virulence of DWV, but could be explained by *Varroa* eliminating other viruses that become more virulent when the mode of transmission changes.

## Results

### Experimental overview

To mimic the effects of changing to a new, vector-based transmission route we serially injected honey bee pupae with viruses and monitored the changes in virus levels. Injecting honeybee extracts into pupae has previously been used to incubate viruses prior to serological experiments^19^ and to obtain standardised inoculum for injection experiments^20^. We adapted this protocol to conduct serial transmission of honeybee extracts by pupal injection for 20+ transmission cycles. We performed two independent transmission experiments with different starting inoculum: (1) extracts obtained from asymptomatic (DWV-naïve) honeybees; and (2) extracts obtained from symptomatic (DWV-infected) honeybees.

### 1. Serial transmission of asymptomatic (DWV-naïive) inoculum

In our first experiment (Figure 1A; Serial Transmission 1), we took our starting inoculum from adults sampled from three asymptomatic honeybee colonies from Sydney, Australia (lacking DWV and naïve to *Varroa*, referred to hereafter as colonies 1, 2 and 3). We subjected white-eyed pupae from the same three colonies to each of three treatments: (1) pupae injected with inoculum containing viruses; (2) pupae injected with extraction buffer as a procedural control (‘buffer’); and (3) pupae left unmanipulated (‘control’). After 4 days, we harvested pupae for extraction to generate inoculum for the next transmission cycle. We passaged inoculum for 20 transmission cycles (18 for colony 3; see Materials and Methods).

**Figure 1:**
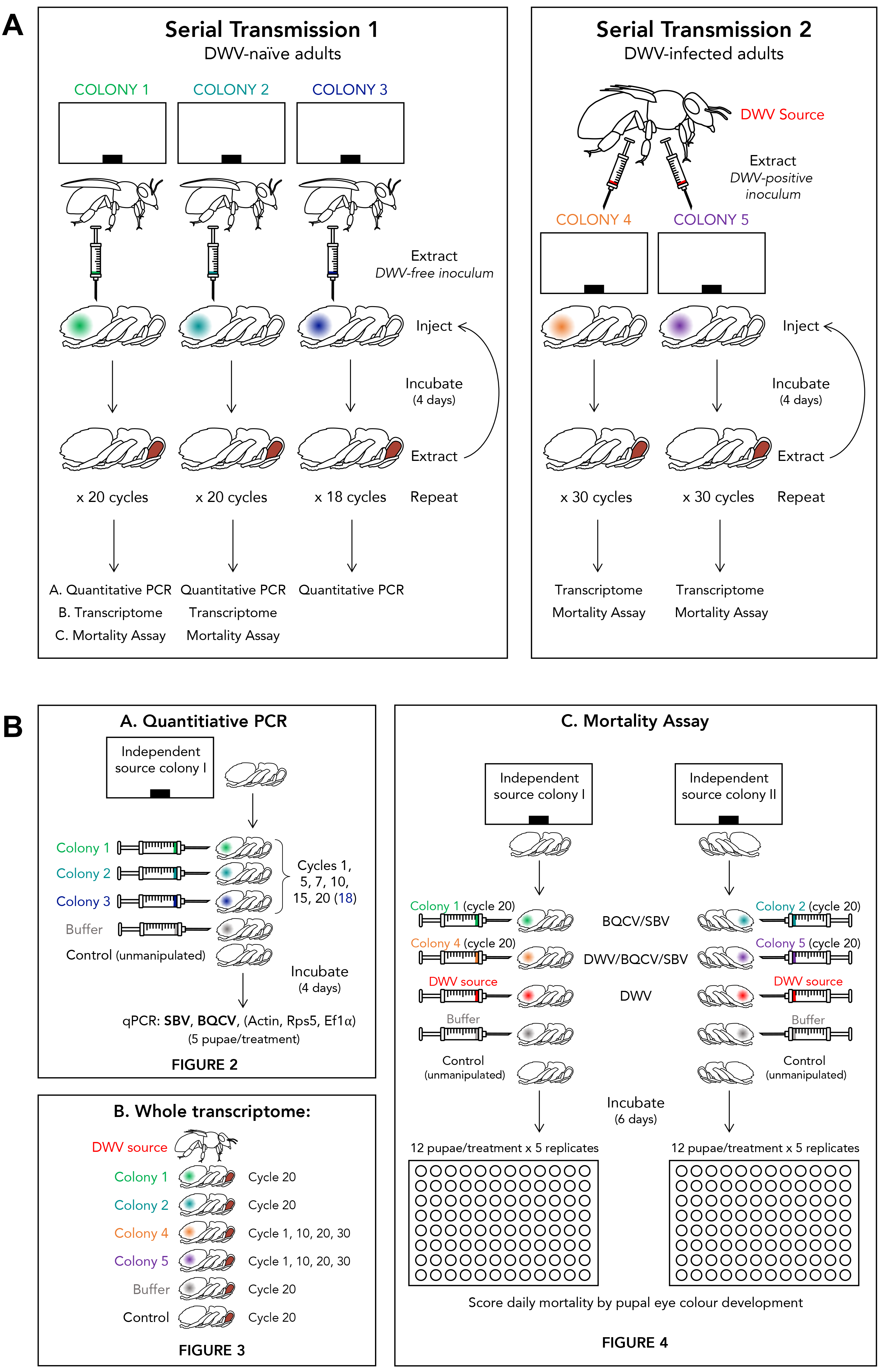
Experimental design of serial transmission experiments. **(A)** Serial transmission 1, with starting inoculum derived from DWV-naïve adults, and injected into pupae from Colonies 1-3 for 20 serial transmission cycles. The number of cycles differed for colony 3 as this colony lost its queen after 18 cycles. Serial transmission 2, with starting inoculum derived from DWV-positive adults from New Zealand, injected into pupae from colonies 4-6. Colony 6 lost its queen early on in the experiment; hence this colony was not included in any further analyses. **(B)** Resulting virus levels and virulence were determined by quantitative PCR (see results in Figure 2), whole transcriptome sequencing (see Figure 3) and mortality assays (see figure 4, and text for further details)

We used end-point PCR to screen for the presence of the five known viruses present in Australia ^21^ in our initial adult workers and in pupae sampled at regular intervals during the 18-20 serial transmission cycles. We detected two just two viruses: Sacbrood virus (SBV) and Black Queen Cell virus (BQCV). Control pupae did not test positive for SBV and BQCV. In contrast, buffer-injected procedural controls occasionally tested positive for SBV and BQCV. It has been well documented that the effect of injection procedure alone can cause the irruption of latent viral diseases in bees^19^, in line with our observations of SBV and BQCV in our buffer, but not unmanipulated, control pupae.

#### Serial transmission results in a rapid increase in viral titre

To determine whether serial transmission resulted in increased viral titres, we assessed expression levels of SBV and BQCV using quantitative PCR and compared those to the expression levels of two endogenous control genes, *Actin* and *Rps5* (see Materials and Methods). We standardised between the three independent colonies and transmission cycles by re-injecting bee extract from colonies 1-3, transmission cycles 1, 5, 7, 10, 15 and 18 (colony 3) or 20 (colonies 1 and 2) into pupae sourced from an independent colony and performed qPCR on these samples, together with buffer-injected and unmanipulated controls.

Both SBV and BQCV virus showed a rapid increase in titre (Figure 2, Table S2). Compared to control and buffer-injected pupae, BQCV levels increased in pupae injected with bee extract after only one transmission cycle, after which levels remained the same (Figure 2 A). Levels of SBV remained low after one transmission cycle but had increased by transmission cycle 5 and remained high thereafter (Figure 2 B).

**Figure 2:**
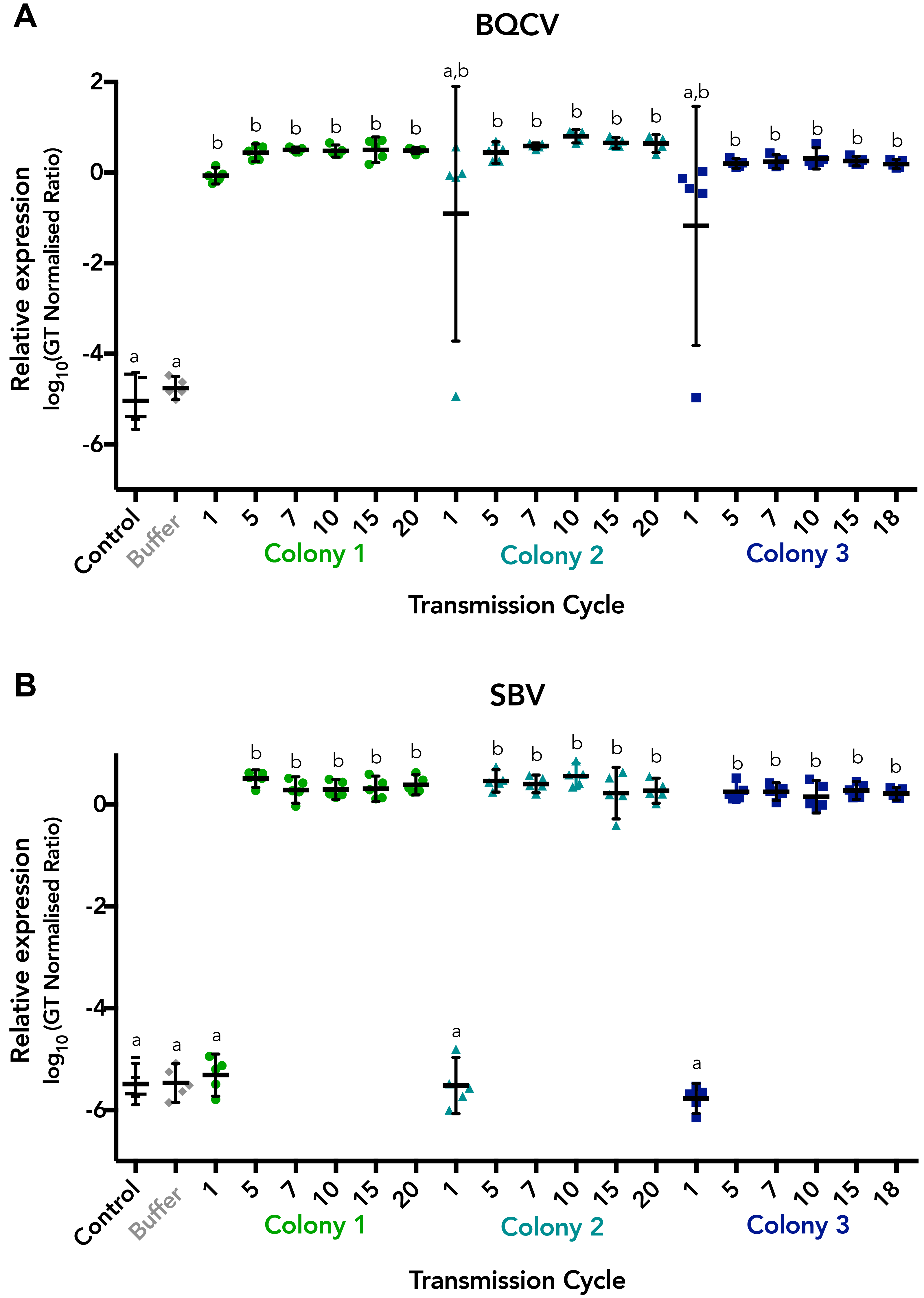
Dot plots showing the relative expression level (log10(GT Normalised Ratio), mean ± 95% CI) of (A) BQCV and (B) SBV compared to two internal honeybee control genes (*Actin* and *Rps5*) in pupae sourced from an independent colony and injected with serially transmitted inoculum from colonies 1-3 (transmission cycles 1, 5, 7, 10, 15 and 18 or 20), and control and buffer injected pupae. Letters indicate which groups differed statistically. See Table S2 for details of the statistical analyses.

To correlate viral titres as measured by qPCR to total RNA content, we examined the amount of viral RNA in pupae injected with bee extract after 20 transmission cycles (colonies 1 and 2, as colony 3 was no longer available due to the loss of the colony’s queen) using HiSeq (Illumina) total RNA sequencing. BQCV and SBV levels made up the vast majority of non-ribosomal RNA in pupae, collectively accounting for 92.6% and 86% of total RNA in colony 1 and 2 pupae, respectively. BQCV levels reached approximately 60%, while SBV levels ranged between 26-35% (Figure 3 A, Table S3).

**Figure 3:**
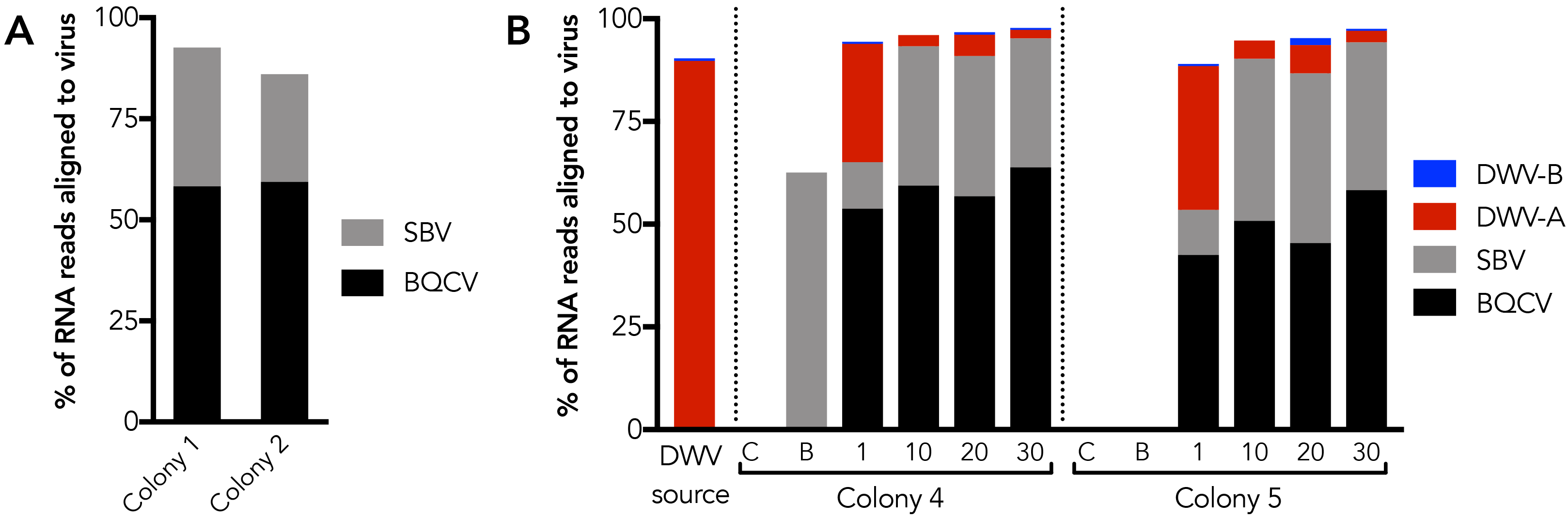
Change in viral titres during (A) serial transmission experiment 1 (DWV-naive); and (B) serial transmission experiment 2 (DWV-positive). A) Levels of SBV (grey) and BQCV (black) in pupae from colony 1 and 2, 4-days post injection with inoculum after 20 serial transmission cycles. Virus levels reached 92 and 86% as a percentage of total RNA, respectively. B) Levels of SBV, BQCV and DWV-strain A (red) and strain-B (blue) in our original inoculum obtained from DWV-positive adults (DWV source), and pupae from colony 4 and 5, 4-days post injection with inoculum after 1, 10 and 20 serial transmission cycles. Also shown are control and buffer pupae from transmission cycle 20. Although our original inoculum (‘Source (DWV)‘) contained exclusively DWV, DWV titres dropped dramatically after injection into pupae, while the titres of SBV and BQCV increased. While our original inoculum mainly contained DWV strain A; the contribution of strain B had increased after 20 transmission cycles, particularly in colony 5, but decreased again after 30 transmission cycles. Data used to produce the figure are presented in Table S3.

### 2. Serial transmission of DWV results in a decrease in DWV titre

We then repeated our serial transmission experiment (Figure 1A, Serial Transmission 2) using inoculum obtained from 5 symptomatic, DWV-infected adult bees from New Zealand (see Material and Methods, including details of quarantine permits), and injecting into lab reared pupae obtained from 2 independent recipient honeybee colonies (referred to as colony 4 and 5; naïve to both DWV and *Varroa*). We passaged inoculum for 30 transmission cycles. We quantified the total amount of viral RNA in the initial adults (DWV source) and after 1, 10, 20 and 30 transmission cycles in pupae 4 days post-injection, along with buffer-injected and control pupae taken from cycle 20 using HiSeq (Illumina) sequencing (Figure 3 B, Table S3).

Almost 90% of non-ribosomal RNA came from DWV in our original inoculum, suggesting that the viral load of symptomatic honey bees can reach extreme levels (Figure 3 B; ‘DWV source’; Table S3). After one transmission cycle, DWV levels reached 25-32% of total non-ribosomal RNA in injected pupae from both colonies. Thereafter, DWV levels decreased rapidly until only a small amount (<10%) of RNA could be attributed to DWV after 10 transmission cycles (Figure 3 B; Table S3). The decrease in DWV titres was accompanied by an increase in BQCV and SBV titre (Figure 3 B), similar to the increase seen in our serial transmission experiment without the inclusion of DWV (Figure 3 A). In the buffer injected pupae from colony 4, we also saw high levels of SBV, indicating that the injection procedure alone can result in increase in endogenous virus levels, in line with previous observations ^19^.

We also saw a shift in DWV strain composition. DWV is known to comprise of 3 main master variants: strain DWV-A, DWV-B and DWV-C^22, 23^. Strain A is globally associated with increased viral titres and colony decline^9, 24^. Strain B is an emerging DWV genotype that has increased virulence compared to DWV-A in laboratory experiments, but has also been found in colonies that seem to cope with the presence of Varroa^17,20,22,23^ (the effect of strain C is currently unknown). Our original inoculum contained low amounts of strain B (0.34% of total viral RNA) which had increased after 20 transmission cycles, particularly in colony 5 (1.66%), only to drop again after 30 cycles (Figure 3 B, Table S3). The total amount of RNA attributable to virus ranged between 88-97% in pupae injected with virus inoculum at all cycles tested, in contrast with control (0.3-0.4% virus) and buffer samples (62% in colony 4 (mentioned above), and 0.18% in colony 5).

### Injecting pupae with serially transmitted SBV and BQCV results in high mortality while DWV alone does not

To compare the virulence of our serially passaged extracts, we injected lab-reared white-eyed pupae with inoculum extracted from our DWV source adults, inoculum from Serial transmission experiment 1, cycle 20 (containing BQCV/SBV, without DWV), and inoculum from Serial transmission experiment 2, cycle 20 (containing DWV/BQCV/SBV). We performed two independent mortality experiments, testing inoculum from cycle 20 from colony 1 and 4 in one independent source colony, and colony 2 and 5 in a second independent source colony (see Figure 1 B for schematic).

Overall survival was significantly affected by treatment in both assays (respectively *χ*^2^4 = 235.68, *p* < 0.00001, *n* = 300 and *χ*^2^4 = 355.21, *p* < 0.00001, *n* = 300; Table S4). Mortality of pupae when injected with DWV alone was not statistically different from buffer-injected controls (both*p* > 0.153; Figure 4, Table S4). When pupae were injected with cycle 20 inoculum from both serial transmission experiments, mortality between inoculum with and without DWV were not statistically different (both*p* > 0.068; Figure 4, Table S4). In both instances, mortality was much higher compared to buffer-injected pupae and pupae injected with DWV alone (all *p* < 0.00001; Figure 4, Table S4). Clearly, increased mortality is due to the increased titres of SBV and BQCV, not due to the presence of DWV. When testing the effect of ‘source colony’ on pupae survival we found that our first source colony had a significantly higher survival than the second (*χ*^2^1 = 4.90, *p* = 0.0268, *n* = 600). However, the overall result was the same for both colonies. ‘Replicate’ had no significant effect on survival in both colonies (respectively *χ*^2^4 = 5.87, *p* = 0.209, *n* = 300 and *χ*^2^4 = 8.84, *p* = 0.065, *n* = 300, Table S4).

**Figure 4:**
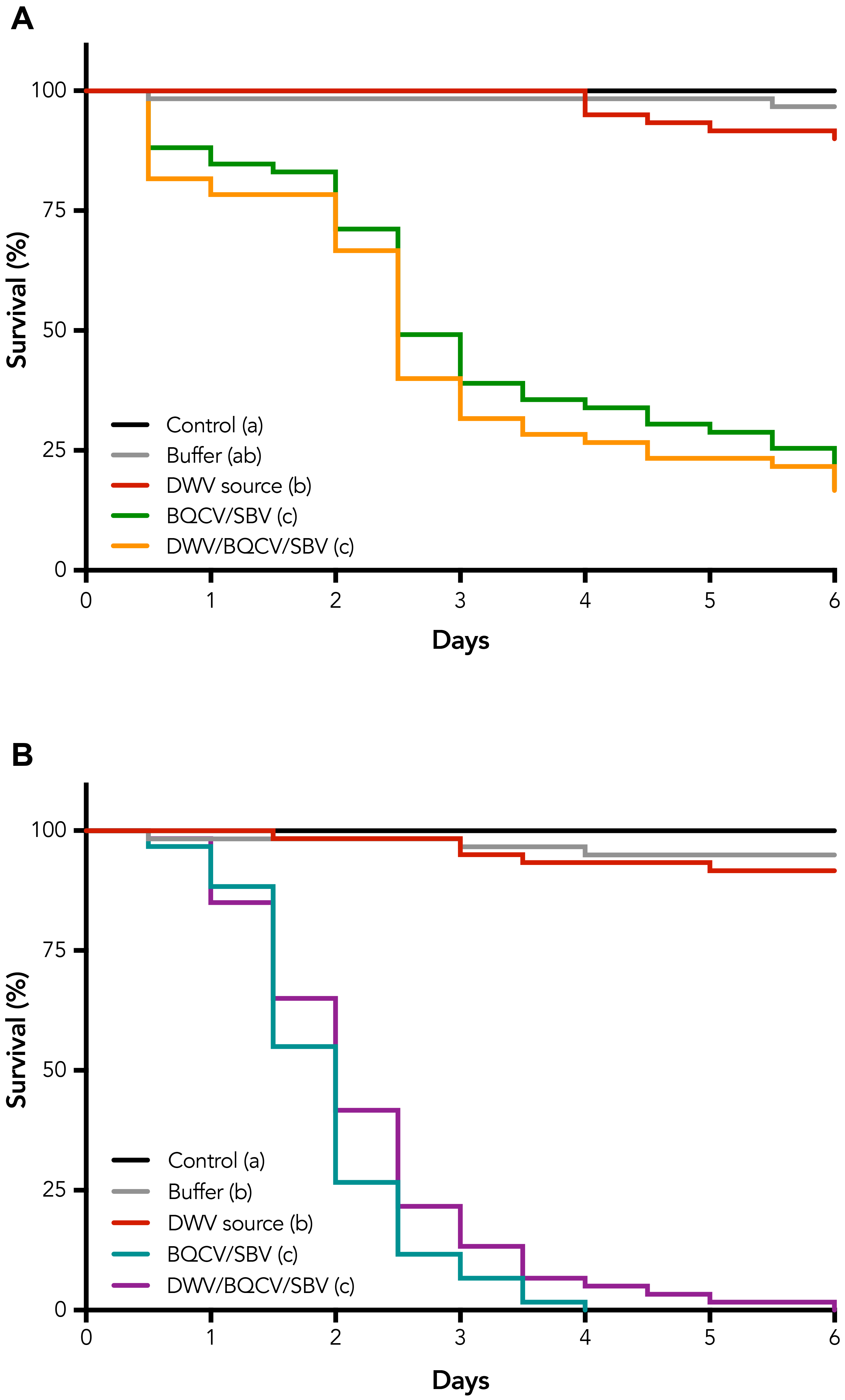
Survival of pupae after injections with inoculum obtained from our original source adults (DWV source) containing mainly DWV strain A, and inoculum extracted after 20 transmission cycles in the absence and presence of DWV, from serial transmission experiment 1 (BQCV/SBV) and serial transmission experiment 3 (DWV/BQCV/SBV). Treatments with the same letter do not differ significantly (see Table 1 for complete statistical analyses). Virus levels present in the different inoculums are given in Figure 3. ‘Control’: unmanipulated pupae; ‘Buffer’: pupae injected with buffer only. (A) BQCV/SBV inoculum from cycle 20 of colony 1; DWV/BQCV/SBV inoculum from cycle 20 of colony 4, injected into independent source colony I; (B) BQCV/SBV inoculum from cycle 20 of colony 2; DWV/BQCV/SBV inoculum from cycle 20 of colony 5, injected into independent source colony II. See Figure S2 for photographs depicting the eye colour change observed in a control and bee extract-injected pupa, used to determine time of mortality in the survival analysis.

## Discussion

We aimed to investigate the effect of changing the route of transmission, from horizontal and, in some cases vertical, to vector-transmitted, to determine if such a change in route of transmission alone is sufficient to increase virulence of RNA viruses contained in the haemolymph of honeybees. We found that two viruses, Sacbrood virus (SBV) and Black queen cell virus (BQCV) rapidly increased in titres when injected into white-eyed pupae. In contrast, when we injected inoculum containing high titres of Deformed wing virus (DWV) strain A, DWV viral titres rapidly decreased, most likely due to competition with SBV and BQCV. Interestingly, injecting high titres of DWV strain A into pupae did not result in the death of the pupae, indicating that this strain of DWV does not kill developing brood. Injecting high titres of SBV and BQCV did result in high mortality.

Both SBV and BQCV are brood diseases; young larvae normally become infected early on via feeding by adult bees^25^. When brood dies from either virus, nurse bees will remove and partially cannibalise the dead brood, thus themselves accumulating the virus. Because both viruses end up in the bees’ hypopharyngeal gland (in which brood food is produced), nurse bees transmit the viruses when feeding young larvae^25^. Under natural conditions, and in the absence of *Varroa*, both SBV and BQCV were found to occur at a frequency of around 10% in summer in Britain using immunodiffusion tests^26^. Both viruses are easily detected when bee extract from adult bees is injected into pupae^27^, indicating that both viruses are present at low incidences without causing overt infections, and readily amplify upon injection into pupae. In Australia, BQCV was found in 65% and SBV in 35% of hives using more sensitive molecular detection methods, further indicating high viral prevalence in the absence of overt infections ^21^. Our results suggest that repeated vector-mediated transmission of bee extract containing SBV or BQCV will rapidly lead to such high viral titres that the brood never develops to adulthood. Our experimental conditions were restricted to pupae, as our quarantine permits required injected pupae to be terminated prior to eclosion. Therefore, our results reflect conditions that are favourable to replication in brood, as we harvested injected pupae randomly, regardless of whether they would have successfully eclosed. Considering that *Varroa* parasitises brood initially, the process of vector-mediated transmission similarly begins in brood. However in contrast to our experimental conditions, only those surviving to eclosion will harbour the viruses that are selected for. This suggests that *Varroa* selects against high replication of viruses causing brood mortality, whereas our selective regime did the opposite.

Another virus commonly found in honey bees, Acute Bee Paralysis Virus (ABPV), cannot replicate when injected into pupae that already contain either SBV or BQCV^26^, showing that indeed SBV and BQCV are highly competitive, probably due to their ability to replicate rapidly. ABPV, and the closely related Kashmir Bee virus (KBV) and Israeli Acute Bee virus (IAPV), are often the first viruses to be associated with the arrival of *Varroa* before they are gradually displaced by DWV^9, 28^. A study documenting the change in viral landscape as *Varroa* invaded the islands of New Zealand, found negative associations between KBV and DWV and between DWV and SBV in both bee and mite samples, while SBV and BQCV were positively associated in both bees and mites^10^. As the time since the arrival of *Varroa* increased, the prevalence of KBV, SBV and BQCV decreased, while DWV increased^10^. These results are consistent with the hypothesis that the succession of honeybee viruses after the arrival of *Varroa* is due to the most virulent viruses being selected against, if *Varroa* transmission facilitates an increase in replication rate ^6, 18^ Our results are the first to provide experimental evidence for this hypothesis.

Clearly honeybee colonies contain a number of different viruses, both of different species as well as different strains of the same species given the high mutation and replication rates of RNA viruses^29^. Competition amongst viruses drives virulence^30, 31^. Inevitably some viruses are more virulent than others. When virulence is too high, the host is likely to die before it has a chance to emerge and transmit the virus to other bees via feeding or faeces. Thus, viruses that are too virulent will be selected against. The arrival of a vector changes the dynamics, as now even highly virulent strains can be transmitted if they manage to get into the vector. But such an increase in the prevalence of highly virulent viruses is bound to be temporary if the vector is killed in the process. Because a vector such as *Varroa* depends on the bee to complete its development (the female mites emerge from the brood cell together with the emerging bee), its arrival will not improve long-term transmissibility of virulent variants, thus leading to the succession from highly virulent viral species to less virulent species as documented in New Zealand^10^.

The last few years have seen a surge in publications that link the arrival of *Varroa* to the emergence of specific strains of DWV^9,10,22–24^. Initially it was thought that DWV strain A was the most virulent strain while strain B was considered to be more benign^22, 32^. However, this simple interpretation now seems questionable, as recently strain B has been associated with colony losses^17^ and appears to be more virulent in an experimental setting^20^. Regardless, the prevailing wisdom is that *Varroa* has led to a change in virulence of an otherwise relatively benign virus by changing the virus’ mode of transmission, thus modifying the virulence-transmission tradeoff^9^. The association between DWV and *Varroa* is so strong, that many now claim that it is the virus that needs to be controlled, not the mite, if we want to protect the bees. We offer experimental evidence for an alternative explanation for the association between *Varroa* and DWV. In the presence of more virulent viruses such as SBV and BQCV, DWV is outcompeted and, if present at all, often below detection level in the absence of *Varroa*. The arrival of *Varroa* quickly selects for an increase in the prevalence of the most virulent viruses until they become so virulent their transmission grinds to a halt due to the death of the brood and thus the mites. Now more benign viruses such as DWV can make their appearance. Hence, perhaps instead of *Varroa* actively selecting for specific, virulent strains of DWV, DWV is simply ‘the last virus standing’ after more virulent species have been selected against.

## Materials and Methods

We used honeybees (*Apis mellifera*) of standard Australian commercial stock that had been kept at the University of Sydney’s apiary for multiple years without showing any symptoms of disease. *Varroa* is not present in Australia. Moreover, it is widely accepted that DWV is not established in Australia after a recent comprehensive, country-wide survey ^21^. A second study showed that strains distantly related to DWV are present in the northern states of Australia (NT and QLD). Next-generation transcriptome sequencing identified contigs showing 53-69% amino acid identity to DWV strains A and B ^33^. This indicates that in some regions of Australia, honeybees host a related virus that is distinct from previously characterised DWV variants. Nevertheless, the colonies used in all of our experiments were sourced from regions where no trace of DWV has been identified in previous surveys, including our own. In addition, we did not detect DWV in our serial transmission experiment or in subsequent next generation sequencing of controls (see further). We thus conclude that our bees were naïve to DWV.

### 1. Serial transmission experiment

#### 1.1 Inoculum preparation

We modified the extraction protocol from Roberts and Anderson^34^. For our first serial transmission experiment, we sampled healthy, DWV-naïve adult bees collected from hive entrances of three separate colonies (colonies 1, 2 and 3, Figure 1A). For each colony, we crushed the thorax and abdomen of five bees in 2ml 0.5 M potassium phosphate buffer (pH 8), removed the lysate by pipetting, then added 5% v/v diethyl ether and 10% v/v chloroform and centrifuged the tubes at 12,000 rpm for 2 minutes. We removed the supernatant and filtered it through a 0.22 μm bacterial filter to remove non-viral pathogens. We then diluted the extracts with potassium phosphate buffer by a factor of 10^−3^. Dilution was necessary because injection of undiluted honeybee extract rapidly kills pupae, potentially due to carryover of toxic metabolites (J. Roberts; personal communication). This dilution factor was chosen based on pilot experiments, where we injected 10-fold serial dilutions of adult and pupal bee extracts into white-eye pupae (ranging from undiluted through to 10^−6^). A 10^−3^ dilution gave the highest concentration that showed no signs of lethality 1 day post-injection. We added 10% v/v green food dye to the bee extract prior to injection to check if injections had been successful. We injected the DWV-free inoculum obtained from each colony into pupae obtained from the same three colonies (1, 2 and 3) as described below.

To obtain DWV for our second serial transmission experiment, we sourced bees visibly showing symptoms of DWV from the top bars of frames from *Varroa* infected colonies in New Zealand (see below for Quarantine details). We cut the thorax and abdomen of five adult bees sagitally in halves and used one half of each bee for inoculum preparation as described above. We kept the other half at −70°C under quarantine conditions for later whole-transcriptome sequencing. We used DWV-containing inoculum as the starting material for injecting into three independent colonies (colonies 4, 5 and 6) as described below.

### 1.2 Injection Procedure

We took 75 white-eyed pupae from their brood comb from each of the six experimental colonies, and distributed pupae into three treatment groups of 25: experimental, buffer and control. We injected the experimental group with 2 μL of initial honeybee inoculum using a Hamilton 10 μl syringe and a 0.3mm needle. As described above, pupae from colonies 1, 2 and 3 were injected with 2μl inoculum taken from DWV-free, asymptomatic nestmate bees, and pupae from colonies 4, 5 and 6 were injected with inoculum obtained from DWV-symptomatic bees from New Zealand (Figure 1 A). We injected the buffer group (procedural control) with 2 μL of potassium phosphate buffer to control for the effect of injection. Experimental and buffer group pupae were injected between the fourth and fifth abdominal tergites. We did not perform any further manipulations on the control group. After injections we placed pupae in petri dishes lined with filter paper soaked in 12% glycerol and incubated them at 34.5°C. All pupae were stored in the lab in our approved quarantine facility under quarantine conditions. After 4 days, we froze the pupae at −70°C until required. To prepare for the next round of injections we selected five pupae for extraction using a random number generator. We randomly selected another five to determine viral levels using real-time quantitative PCR, and kept the remaining fifteen in reserve. Extracts from previous transmission cycles were then injected into the next round of white-eyed pupae collected from brood combs originating from the same six experimental colonies. For the DWV-naïve transmission experiment, we concluded a total of 20 transmission cycles for colonies 1 and 2 whereas the third colony replaced its queen so that we were unable to collect pupae beyond 18 transmission cycles. For our DWV-positive transmission experiment, we concluded 30 transmission cycles for colonies 4 and 5, whereas the queen from colony 6 was replaced during the 4^th^ transmission cycle and thus this colony was excluded from any further analysis.

### *1.3* Detection of viruses

We used end-point PCR to screen for the presence of viruses in our starting colonies. We used Trizol (Life Technologies) to extract RNA from 12 uninjected bees collected from colonies 1-3 at the beginning of the first serial transmission experiment. For our second serial transmission experiment we extracted RNA from 6 pupae from colonies 4-6, sampled at the time of transmission cycle 1. We quantified RNA using a Qubit Broad Range Assay (Life Technologies), and normalised to 200 ng/μl before treatment with DNAse. We synthesised first strand cDNA from 0.5 μg total RNA using SuperScript III Reverse Transcriptase (Invitrogen) and random hexamer primers. We performed PCRs to screen for presence of BQCV, SBV, IAPV and Lake Sinai Virus (LSV), as these viruses are most commonly found in Australian bees ^21^, using the primers described in Table S1 with an initial 5 min denaturing step at 94 °C, followed by 38 cycles of 94 °C for 1 min, annealing temperature for 1 min and 72 °C for 1 min per kb of product, with a final extension step for 10 min at 72 °C. We visualised PCR products on a 1.5% agarose gel using SYBR Safe DNA stain (Life Technologies). Positive PCR products were sequenced by Macrogen and identity confirmed by BLAST to NCBI GenBank online database.

We screened for viruses throughout the experiment using endpoint PCR at various timepoints. For serial transmission experiment 1, we examined 5 pupae from all three treatment groups at transmission cycles 1, 5, 7, 10, 13 and 15. For serial transmission experiment 2 we examined 3 pupae from all treatment groups at transmission cycles 1, 10, and 20 to validate the presence of DWV in our experimental group, and the absence of DWV from our buffer and control groups.

### 2. Assessment of virus levels

#### 2.1 Serial transmission experiment 1- Quantitative Real-time PCR

To compare the viral titres between colonies 1-3 after serial transmission with extracts sourced from DWV-naïve bees, we collected white-eyed pupae from an independent colony to standardise for colony background. We injected bee extracts from experimental groups of colonies 1-3, from transmission cycles 1, 5, 7, 10, 15 and 20 (cycle 18 for colony 3), into 10 white-eyed pupae, along with 10 buffer-injected procedural controls and 10 uninjected (unmanipulated) controls (see Figure 1 B for a schematic representation of the experiment). We randomly selected five pupae from each group and extracted RNA from each pupa separately in 1 mL of Trizol. We treated the RNA with DNAse and performed cDNA synthesis using the same method described above. We then diluted cDNA to a final concentration of 27 ng/μl. We created negative controls for the qPCR assay by pooling extracted RNA from samples drawn from the same treatment group and transmission cycle, treating them with DNAse and mixing them with all the reagents for cDNA synthesis except the reverse transcriptase enzyme. We created the standards for our qPCR assay by taking previous cDNA samples with high levels of SBV and BQCV, then performing a serial dilution over 3 orders of magnitude.

We designed qPCR primers to amplify SBV and BQCV and used previously published primers for β-Actin, Ef1-α, and Rps5, which served as endogenous controls (Table S1). We confirmed the specificity of each primer pair via melt-curve analysis and gel electrophoresis. We performed the assay using a Roche LightCycler 480 using 2x SYBR Master Mix (Roche Technologies). We pre-incubated the reactions (95°C, 10 minutes) prior to 45 amplification cycles (95°C, 10 seconds; 58°C 10 seconds; 72°C, 10 seconds), and measured fluorescence at each extension step. We obtained Cq values using the second derivative maximum method using the Roche LightCycler 480 software. The same software was used to calculate the efficiencies of each set of primers from the standard curves on each plate.

##### Statistical analysis - quantitative real-time PCR

We compared the stability of each reference gene in Bestkeeper^35^ and used the two most stable reference genes, *Actin* and *Rps5*, to normalise the expression of BQCV and SBV in all samples. The expression level of each gene was calculated as 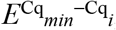, where E is the efficiency of the primers calculated, Cq*min* is the lowest Cq value for a given gene and Cq*i* is the Cq of sample *i*. The expression of each of SBV and BQCV in each sample was then normalised against the geometric mean of the expression levels for both reference genes to obtain the final relative expression score^36^. To compare the viral titres between each transmission cycle and between colonies 1-3 in our pupal quantitative PCR, we performed a one-way ANOVA (Table S2).

#### 2.2 Serial transmission experiment 2- Whole transcriptome RNA sequencing

To compare virus levels between colonies 4 and 5 after serial transmission with DWV-inoculum, we extracted RNA from five randomly selected pupae 4 days post-injection for each of transmission cycles 1, 10, 20 and 30, as well as buffer injected and unmanipulated control pupae from transmission cycle 20. We also extracted RNA from five pupae from transmission cycle 20 from colonies 1 and 2 from serial transmission experiment 1 (DWV-naïve transmission). Finally, we extracted RNA from our DWV source population, using the stored half of adult bees from which the haemolymph containing DWV was initially extracted (see Figure 1 B for schematic). We extracted RNA using 1mL Trizol as outlined above. After DNase treatment, we used an RNeasy Mini Kit (Qiagen) to clean RNA to a total of ≥5μg. Cleaned RNA was diluted to a minimum of 100ng/μL into a 50μL aliquot and stored at −70°C until it was shipped on dry ice to the Australian Genome Research Facility (AGRF) laboratory (Melbourne, Australia) for preparation of whole transcriptome, 100bp paired-end library with ribosome depletion (Ribo-zero Gold (human/mouse/rat)), and HiSeq (Illumina) sequencing. We multiplexed fifteen libraries across two lanes, providing between 3.13-4.2 Gb data per library, for a total data yield of 56.74 Gb. The raw sequencing reads from this project have been deposited to Genbank under the Bioproject ID PRJNA397460 at the Sequence Read Archive (SRA Study ID: SRP114989).

##### Sequencing - data analysis

We performed an intial *de novo* assembly of sequencing reads for each sample using Trinity^37^. To determine which honey bee viruses were present, we used BLAST searches to compare Trinity-assembled contigs to a custom honey bee virus database containing all currently known honey bee virus genome sequences. We found contigs matching to BQCV and SBV in all samples. DWV contigs were present in assemblies from the DWV source population and cycles 1-30 of the DWV serial transmission experiment. In addition, the DWV source population also contained contigs matching to the recently described Apis Rhabdovirus 1 and 2 (ARV-1 and ARV-2)^38^. However the levels of these viruses were below 0.05% of the total RNA reads (Table S3), and were subsequently not detected in any further transmission cycles. We found no other viruses in our samples. Interestingly, short contigs for DWV were also assembled from our buffer injected and unmanipulated controls. Previous studies using Illumina HiSeq and MiSeq technology have reported ‘sample bleeding’ due to reads being incorrectly assigned to the wrong sample source when multiplexed in the same sequencing lane^20, 39^. To assess the level of DWV read misassignment in our buffer and control samples, we aligned sequencing reads of each sample to DWV-A and DWV-B genomes using Bowtie2^40^. All control samples showed less than 0.02% of total reads aligning to DWV. This level was similar to the level of multiplex sample bleeding that we observed in a negative control sample. In addition, we were unable to amplify DWV using PCR from cDNA synthesised from our buffer and control samples, therefore we concluded that the DWV contigs present in our controls are a result of inaccurate sample assignment of reads during the multiplexed HiSeq sequencing run.

Prior to analysing the viral content of our transcriptomes, we assessed the proportion of residual ribosomal RNA (rRNA) reads, as complete ribosome depletion may not be obtained when using Ribo Zero Gold (human/mouse/rat) for invertebrate samples. We identified *Apis mellifera* rRNA from Trinity contigs by BLAST searches and aligned reads for each sample using Bowtie2. The percentage of residual rRNA reads ranged from 2.5 - 46% of total RNA reads per library (Table S3). These values were factored into any subsequent viral percentage calculations.

To determine the viral content in each treatment condition, we aligned sequencing reads of each sample to BQCV, SBV, DWV-A and DWV-B genomes using Bowtie2. We used representative SBV and BQCV contigs assembled *de novo* from our samples as the template for Bowtie2 alignments, as our SBV and BQCV strains differed significantly to the reference SBV and BQCV genomes from Genbank (SBV: 92.6% nucleotide identity to AF092924^41^; BQCV: 89.2% nucleotide identity to AF183905^42^). We used DWV-A and B reference sequences available in Genbank (AJ489744 and AY251269) as the template for Bowtie2 alignments, as the DWV strain assembled from our DWV source population matched to DWV-A with high nucleotide identity (98.7%). The reference SBV, BQCV and source DWV sequences used in this study have been deposited to Genbank under accession numbers MF623170, MF623171 and MF623172. We compared the SBV and BQCV sequences from our study to published available SBV and BQCV genomes. We performed nucleotide alignments in Geneious (v10.2.4 ^43^) using Muscle and generated phylogenetic trees using PhyML (Figure S1).

#### 3. Assessment of virulence

##### 3.1 Pupal survival screen

To assess the virulence of the viruses contained in the inoculum generated after serial transmission cycles in experiment 1 (containing SBV/BQCV) and experiment 2 (containing DWV/SBV/BQCV), we developed a mortality assay using change in pupal eye colour to determine pupal mortality. As pupae develop, pigments such as ommochromin are deposited in the compound eyes and ocelli, causing a change in colour from white, through pink and red, to the endpoint black^44^ (see Figure S2). By comparing the colour of a pupa’s eyes over consecutive days, we developed an assay that allowed us to determine the point in time a pupa died. A pupa was determined to have died when its eyes had ceased changing colour over two consecutive photographs, and/or when the compound eye had retracted from the cuticle (Figure S2).

We sourced pupae from 2 independent honeybee colonies and injected them with inoculum as per our serial transmission protocol. We injected inoculum from transmission cycle 20 from Colony 1 (BQCV/SBV) and Colony 4 (DWV/BQCV/SBV) for the first trial, and Colony 2 and Colony 5 for the second trial, along with the DWV source inoculum, buffer and unmanipulated controls (12 pupae per replicate, 5 replicates per colony, see Figure 1 B for schematic). We placed pupae into 0.6mL 96 well PCR plates so that we could monitor their development by taking photographs of their eyes two times per day. We used a 1.5% w/v agar gel as a substrate to maintain moisture and standardise the height of pupae, and added 0.01g 100mL^−1^ copper sulphate to the hot agar prior to pouring in order to inhibit fungal growth. We photographed pupae using a Nikon D5100 camera with a Tamron 60mm F/2 macro lens and terminated the experiment at day 6 prior to eclosion.

###### 3.1 Statistical analysis - survival screen

We compared the pupae’s survival with Cox’s proportional hazards survival analyses using R-3.3.3^45^ with the package survival^46^. We checked the log-linearity of covariates by plotting the Cox models’ martingale residuals against fitted values^47^. We checked the proportional hazards assumption of the Cox regression models following Grambsh & Therneau^48^ (all *p* > 0.05). In each Cox model we investigated for each colony the effect of ‘treatment’ (i.e. control, buffer, pure DWV, serially transmitted BQCV and SBV, serially transmitted BQCV, SBV and DWV) on pupae survival. We also included ‘replicate’ as a covariate, as well as the interaction between treatment and replicate in all models. Since this interaction proved to be non significant in both models (respectively colony 1: *p* = 0.214 and colony 2: *p* = 0.244), we removed the interaction and recalculated the model. Post-hoc p-values were corrected for multiple comparisons following the Benjamini and Hochberg procedure ^49^.

###### Quarantine permits

Frozen worker honey bee samples containing Deformed wing virus were imported from the New Zealand Institute for Plant and Food Research, Hamilton, under our Department of Agriculture and Water Resources import permit 0000917783. To work with DWV in Australia, we require a quarantine permit that restricts us from injecting imported viruses into adult bees. However, we are permitted to inject into pupae provided we terminate experiments prior to eclosion. Therefore, we injected inoculate into white-eyed pupae, and we terminated each cycle after 4 days to enable sufficient time for viral replication, while avoiding eclosion (Figure 1 A).

###### Data Availability

The raw sequencing reads from this project have been deposited to Genbank under the Bioproject ID PRJNA397460 at the Sequence Read Archive (SRA Study ID: SRP114989). The reference SBV, BQCV and source DWV sequences used in this study have been deposited to Genbank under accession numbers MF623170, MF623171 and MF623172.

## Acknowledgements

We thank the Australian Research Council (ARC) for financial support (FT120100120 and
DP170100844 to MB) and the University of Sydney’s Marie Bashir Institute for Infectious Diseases and Biosecurity for seed funding (to EJR and MB). BY is supported by the Fyssen
Foundation. We thank The New Zealand Institute for Plant & Food Research for providing DWV samples.

## Author contributions

EJR and MB designed the experiments. EJR, NM, TLG and BY performed the experiments.

EJR, NM, TLG, BY and MB analysed the data. EJR and MB wrote the paper.

